# FGF activity asymmetrically regulates the timing of habenular neurogenesis in a Nodal-dependent manner

**DOI:** 10.1101/261834

**Authors:** Benjamin J. Dean, Joshua T. Gamse, Shu-Yu Wu

**Affiliations:** Department of Biological Sciences, Vanderbilt University, Nashville, TN 37232, USA; Current address: Division of Pediatric Neurology, Seattle Children’s Hospital, Seattle, WA 98105; Current address: Department of Reproductive Toxicology, Drug Safety Evaluation, Bristol-Myers Squibb, New Brunswick, New Jersey.; Current address: Department of Molecular Physiology and Biophysics, Vanderbilt University, Nashville, TN 37232, USA

## Abstract

The highly conserved habenular nuclei in the vertebrate epithalamus function as an integrating center that relaying information between the forebrain and the brain stem. These nuclei play crucial roles in modulating a broad variety of cognitive behaviors. Moreover, habenular nuclei has also attracted interest as a model for brain asymmetry, since many vertebrates exhibit left-right differences in habenular size and neural circuitry. Left-right (L/R) asymmetry is a shared feature of the central nervous system in vertebrates. Despite its prevalence and functional significance, few studies have addressed the molecular bases for the generation of the asymmetric brain structure, perhaps due to the absence of genetically accessible model animals showing robust brain asymmetry. Previous studies on zebrafish epithalamus demonstrated that Nodal signaling directs the habenular asymmetry during the early stages of development by biasing the neurogenesis on the left-side. Here, we discover a novel regulatory module involving asymmetric activation of FGF signaling that determines the timing of habenular neurogenesis by regulating cell-cycle progression of neuronal progenitors, which seamlessly integrates the L/R patterning driven by Nodal and the spatiotemporal patterning of habenular neurons.

## INTRODUCTION

Vertebrate brain function depends on generating a great variety of neuronal subtypes during development. Excitatory, inhibitory and neuromodulatory neurons must be generated in carefully balanced numbers. Errors in neuronal specification play a key role in the pathophysiology of many neurodevelopmental disease states including epilepsy, autism and schizophrenia (Levitt et al., 2004; Rubenstein, 2010).

All neurons of the central nervous system arise from a pseudostratified neuroepithelium (NE) composed of neural progenitors (NPs). NPs get spatial and temporal information from their dynamic exposure to secreted morphogens and their inhibitors, including members of the FGF, TGFβ, Wnt, Bmp, Shh and Retinoic acid families. One or more morphogenic signaling pathways are then integrated and drive expression of specific sets of homeodomain-containing and basic helix-loop-helix transcriptions factors (Guillemot, 2007). From this combinatorial code NPs become specified and go on to generate a restricted repertoire of neurons at specific developmental stages.

NPs divide in either proliferative or neurogenic divisions. In proliferative divisions both daughters retain progenitor status while in neurogenic divisions at least one daughter exits the cell cycle and terminally differentiates. Over time, as NPs are specified, they transition away from purely proliferative divisions to neurogenic divisions. The timing and execution of this ‘neurogenic switch’ and subsequent differentiation are regulated by extrinsic factors and are fundamental to generating the proper number and subtype of neuronal daughters (Aizawa et al., 2007; Scholpp et al., 2009).

Most of our understanding of spatiotemporal patterning and neurogenesis has resulted from careful exploration of powerful model CNS regions such as the dorsoventral (D/V) axis of the spinal cord and anteroposterior (A/P) axis of the thalamus (Kiecker and Lumsden, 2012; Scholpp et al., 2009). These, and other studies, have shown the importance of both spatial as well as temporal cues in determining proper neuronal cell fates along the A/P and D/V body axis. However, comparatively little is known about how the timing of neurogenesis effects cell fate across the left-right (L/R) axis.

L/R brain asymmetry is highly conserved across vertebrates and is crucial for normal behavior. The general mechanism of spatiotemporal neuronal patterning must be integrated with L/R patterning at some level to give rise to the neuronal substrates of asymmetric brain function (Bianco and Wilson, 2009). The dorsal habenular nuclei (dHb) provide a rich model to determine the pathways by which neurogenesis and L/R patterning are integrated. These bilaterally paired brain structures lie in the dorsal diencephalon, are embedded in monoaminergic circuitry and have some of the highest concentrations of acetylcholine and GABA receptors in the entire brain (Viswanath et al., 2013). In many classes across the vertebrate lineage these nuclei are asymmetric (Bianco and Wilson, 2009). In the zebrafish, *Danio rerio*, the dHb contain medial and lateral subnuclei. The subnuclei show unique patterns of genes expression and are asymmetric in neuronal fate allocation across the L/R axis. It has been previously reported that habenular neurogenesis begins asymmetrically and that this asymmetry is dependent on the left-fate-determining Nodal signaling pathway (Regan et al., 2009). It has also been observed that the timing of neurogenesis for a particular neuron correlates with its fate as a medial or lateral habenular neuron (Aizawa et al., 2007). To date it is unclear how Nodal ligands direct the asymmetric timing of habenular neurogenesis and ensure properly asymmetric distribution of neuronal subtypes in the dorsal habenula. Here we report that FGF signaling serves as a point of integration of Nodal-mediated L/R patterning and habenular neurogenesis.

The FGF signaling pathway plays a crucial role in patterning the vertebrate brain. *Fgf8* is robustly expressed in the dorsal diencephalon (DD) during habenular development. Indeed, loss of *fgf8* signaling leads to a failure to form the habenular nuclei in the mouse and hypomorphic alleles of *fgf8* in the zebrafish result in defects in habenular development (Martinez-Ferre and Martinez, 2009; Regan et al., 2009). FGF signaling, and *fgf8* specifically, are known to regulate the neurogenic switch in NPs near the midbrain-hindbrain boundary (Lahti et al., 2010; Saarimäki-Vire et al., 2007). In particular, high levels of FGF signaling maintain proliferative divisions, but reductions in FGF signaling allow neurogenic divisions. These studies support a classical view of FGF as a dose-dependent developmental effector (Garcia-Maya et al., 2006). Thus, we hypothesized that the timing of habenular neurogenesis would similarly depend on levels of FGF signaling. Taking advantage of small molecules, we tune FGF signaling levels up and down and demonstrate that FGF regulates the timing of habenular neurogenesis in a dose-dependent manner and furthermore, that FGF exerts its effects by regulating the cell-cycle dependent kinase inhibitor (CDKI), *kip2*. Interestingly, perturbations in the timing of dorsal habenular neurogenesis have downstream effects on neuronal fate Finally, we find that FGF-regulated neurogenesis is rendered asymmetric by Nodal-driven inhibition of FGF signaling activity in the left habenula. To our knowledge this is the first report of Nodal regulating the FGF pathway and propose that FGF signaling serves as a key regulator of asymmetric habenular neurogenesis integrating L/R and neurogenic patterning.

## MATERIALS AND METHODS

### Zebrafish maintenance and strains

Zebrafish were raised at 28.5°C on a 14/10 hour light/dark cycle and staged according to hours post-fertilization. The following fish lines were used: the wild-type strain AB*\** (Walker, 1999), *Tg(-8.4neurog1:GFP)* (Blader et al., 2003), *TgBAC[dbx1b:eGFP]* (Koyama and Kinkhabwala, 2011), *TgBAC[dusp6:d2eGFP]* (Molina et al., 2007), *fgf8^x15^* (Kwon and Riley, 2009), *tbx2b^c144^* (Snelson et al., 2008), *flh^n1^* (Talbot et al., 1995). All experiments were approved by the Vanderbilt University’s Institutional Animal Care and Use Committee (IACUC) and Office of Animal Welfare, and performed according to national regulatory standards.

### Whole mount *in situ* hybridization

Whole-mount RNA *in situ* hybridization was performed as described previously (Gamse, 2003), with one change: 5% dextran sulfate was added to the hybridization buffer. Hybridized probes were detected using alkaline phosphatase conjugated antibodies (Roche) and visualized by 4-nitro blue tetrazolium (NBT; Roche) and 5-bromo-4-chloro-3-indolyl-phosphate (BCIP; Roche) staining for single colorometric labeling, or by NBT/BCIP followed by iodonitrotetrazolium (INT) and BCIP staining for double colorometric labeling.

### Whole mount fluorescent *in situ* hybridization and immunohistochemistry

Whole-mount fluorescent *in situ* hybridization and immunohistochemical co-labeling was performed as described previously (Doll et al., 2011), with the following additional reagents: 5% dextran sulfate was added to the hybridization buffer. Hybridized probes were detected using anti-DIG alkaline phosphatase conjugated antibodies (Roche). In addition to Fast Red substrate (Sigma, F4648), some experiments used 3 x 5 minutes washes in Fast Blue Buffer (Lauter et al., 2011) and were developed in Fast Blue Substrate (0.25mg/mL Fast Blue Substrate and 0.25mg/mL nAMP in Fast Blue Buffer) diluted in Fast Blue Buffer. In addition to the anti-DIG antibody, the primary antibodies used were mouse anti-HuC (1:400, Life Technologies), rabbit anti-GFP (1:500, Torrey Pines Biolab), chicken anti-GFP (1:300, Vanderbilt Antibody and Protein Resource), rabbit anti-Kctd12.1 (1:300, Gamse et al. 2005). Primary antibody was detected using goat-anti-rabbit, goat-anti-mouse or goat-anti-chicken antibodies conjugated to Alexa 488, Alexa 568 or Alexa 633 fluorophores (1:300, Molecular Probes).

Double fluorescent in situ hybridization was performed with the following modifications to the above colorometric *in situ* protocol. After hybridization of DIG and fluorescein labeled probes, anti-DIG antibody was applied (1:5000, Roche) overnight at 4°C. The following day embryos were washed 4 x 20 min in PBS with Triton (PBSTr) and 3 x 5 min in Fast Blue Buffer and developed in Fast Blue Substrate diluted in Fast Blue Buffer. After color development, embryos were washed 2 x 10 min in PBSTr. The alkaline phosphatase was acid inactivated by a 10 min wash in 0.1M glycine HCl pH2.0. After 2 x 10 min PBSTr washes, embryos were incubated in anti-fluorescein antibody (1:1000, Roche) overnight at 4C. The following day, color was developed in Fast Red substrate as in Doll et al. 2011.

*dbx1b* probe (Gribble et al., 2007) was produced from pCRII-*dbx1b* plasmid linearized by BamHI and transcribed by T7 RNA polymerase. *her6* probe (Scholpp et al., 2009) using pBSSK-*kip2*, NotI and T7 RNA polymerase, *kip2* probe using pBS+-*kip2*, NotI and T7 RNA polymerase and *lefty1* probe using pBSSKII-*lefty1* with NotI and T7 polymerase.

### Inhibitor treatments

For whole-mount *in situ* hybridizations and antibody labeling, embryos were incubated in their chorions in 1uM of SU5402 (Tocris), 10uM, 25uM or 50uM of BCI (Sigma) or 50uM SB505124 dissolved in 0.3% dimethyl sulfoxide (DMSO) in egg water supplemented with 0.003% N-phenylthiourea (PTU; Sigma-Aldrich) to prevent melanin formation. Control embryos were treated with 0.3% DMSO in parallel with their SU5402-treated, BCI-treated or SB505124-treated siblings. SU5402 and BCI treatments were all from 25-26hpf unless otherwise stated. Embryos were either fixed immediately following treatment or SU5402/BCI/DMSO was washed off with 5 x 5 min egg water before being returned to egg water with PTU to develop to the desired stage for fixation. SB505124 treatments were from 16hpf until fixation.

### Imaging and Microscopy

For fixed tissue, samples were cleared in a glycerol series (50%, 100%). For in-vivo time-lapse microscopy, embryos were anesthetized in 1% Tricaine and mounted in 0.6% agarose containing 0.04% Tricaine and 0.003% N-phenylthiourea (PTU; Sigma-Aldrich) to prevent melanin formation. Time-lapse images were collected every 15 minutes for the hours indicated on a Zeiss/Perkin Elmer spinning disk confocal microscope. Fixed-tissue fluorescent images were collected on the same Zeiss/Perkin Elmer or a Zeiss LSM 510 Meta confocal microscope. Time-lapse and fixed tissue images were taken with a 40X oil-immersion objective and analyzed with Volocity software (Improvision).

### Quantitation and Statistics

Quantitation of wildtype *her6* and *kip2* expression levels took advantage of the *TgBAC(dbx1b:GFP)* transgenic line which marks the entire habenula. Using Volocity software, the left or right habenula was selected using the GFP channel, and then total fluorescence was recorded from the *kip2* or *her6* channel. Because FGF regulates *dbx1b* this approach was not valid in *fgf8^x15/x15^* mutants. To quantitate relative fluorescence levels of *Tg(dusp6:d2eGFP)* as well as *her6* and *kip2* expression levels in the *fgf8^x15/x15^* background, three optical sections were taken through the habenula. Using the ventral margin of the pineal gland as an anchor, sections 5um dorsal, 5um ventral and through this anchor were selected allowing for uniformity across samples. Then the fluorescence of the entire left or right diencephalon was measured in each optical section using Volocity (Improvision). These values were then totaled and taken as representative of the habenular expression. To compare expression levels on the left and right, or to compare the ratio of expression between two groups, Student’s T-test was performed. To measure asymmetry of neurogenesis in SU5402 and BCI-treated embryos we first counted GFP+ or HuC+ neurons and subsequently employed a Wilcoxon signed-rank test (Roussigné et al., 2009).

## RESULTS

### FGF signaling is asymmetric during early habenular development

Previous analysis of the DD of zebrafish *fgf8^ti282/ti282^* hypomorphs showed a probable role for FGF signaling in habenular development (Regan et al., 2009). Subsequent analysis of *fgf8^x15/x15^* null mutants confirmed that FGF signaling is crucial for normal habenular development. In the absence of *fgf8*, the developing habenula have decreased proliferation, increased apoptosis and the cells that are produced fail to differentiate in neurons (Supplemental Figure 1A-C). Thus, FGF signaling regulates the number of habenular cells, their survival and is required for their proper differentiation. However, the severity of the null mutant makes is hard to separate how FGF signaling regulates neurogenesis versus differentiation.

To understand habenular FGF signaling, we took advantage of a validated FGF reporter line where a *dusp6* promoter element drives expression of a destabilized enhanced-GFP (Molina et al., 2007). *dusp6* is a member of the dual-specificity phosphatase family of phosphatases and is a feedback inhibitor of the FGF signaling pathway. Thus, this reporter is activated in the presence of robust FGF signaling. The enhanced-GFP used is fused to a PEST sequence leaving the mature protein with only a 2 hour half-life (Li, 1998). As a transgenic line, *TgBAC(dusp6:d2eGFP)* allow for analysis of FGF signaling activity in fixed tissue as well as in vivo time-lapse confocal microscopy revealing dynamic changes in FGF signaling activity.

Strikingly, a robust asymmetry in habenular FGF activity was evident at 26hpf, at the beginning of habenulogenesis (Figure 1A&B). FGF activity was higher in the right habenula and lower in the left habenula (Figure 1B). To confirm that asymmetric FGF activity was in the habenulae, we analyzed the co-expression of this transgene with the recently reported dHb progenitor marker, *dbx1b* (Dean et al., 2014). Single optical sections reveal that asymmetric FGF signaling is present in the early habenula (Figure 1C&D).

**Figure 1:**
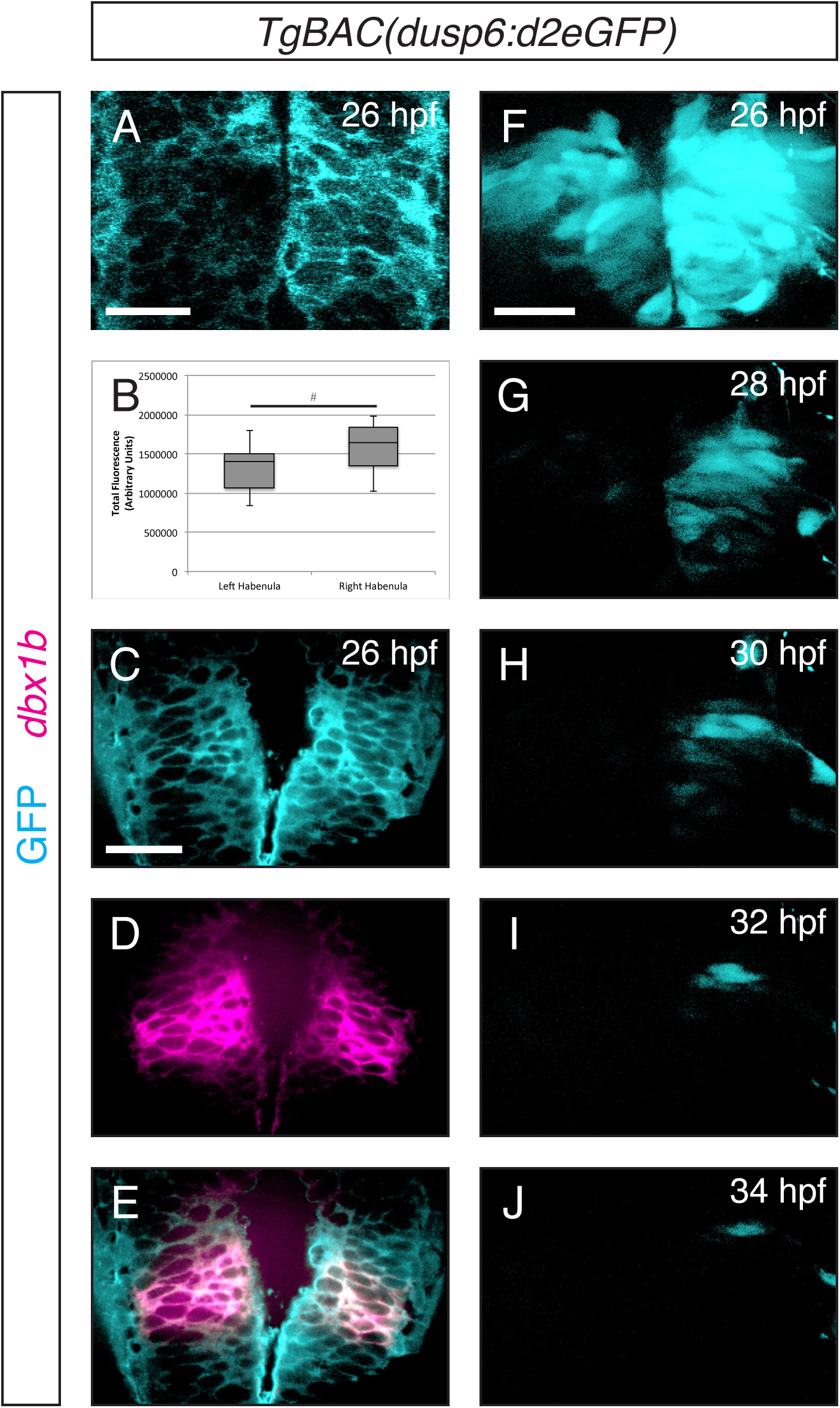
FGF signaling is asymmetric in the early developing habenulae. The FGF reporter *TgBAC(dusp6:d2eGFP)* has higher levels of expression in the right habenula than the left at 26hpf (A-B). This expression pattern co-localized with the early habenulae markers *dbx1b* (C-E). Using in vivo time-lapse confocal microscopy we observed that FGF signaling decreases between 26 and 34hpf falling below detectable levels in the left habenula (28hpf) before loss of signal on the right (34hpf, F-J). ^#^p<1.8 x 10–^6^. Scale bars are 50uM.

Taking advantage of the destabilized eGFP in our reporter, we employed in-vivo time-lapse confocal microscopy to study the temporal dynamics of FGF signaling in the habenula. Between 26hpf and 34hpf asymmetry in transgene expression persists while overall levels of signaling decline on both sides reaching undetectable levels at 28hpf and 32hpf respectively (Figure 1F-J). We concluded that in the early stages of habenular development, FGF signaling is greater in the right habenula. As development proceeds, FGF signaling activity decreases greatly, but in a stable asymmetric fashion.

### Asymmetric habenular neurogenesis is FGF-dependent

It has been previously reported that post-mitotic neurons appear in the habenula asymmetrically, with the earliest born neurons appearing in the left habenula about 4 hours before they appear on the right (Roussigné et al., 2009). Neurogenesis begins shortly before 36hpf and the asymmetric distribution of neurons is apparent through 40hpf (Figure 2A, C and I).

**Figure 2:**
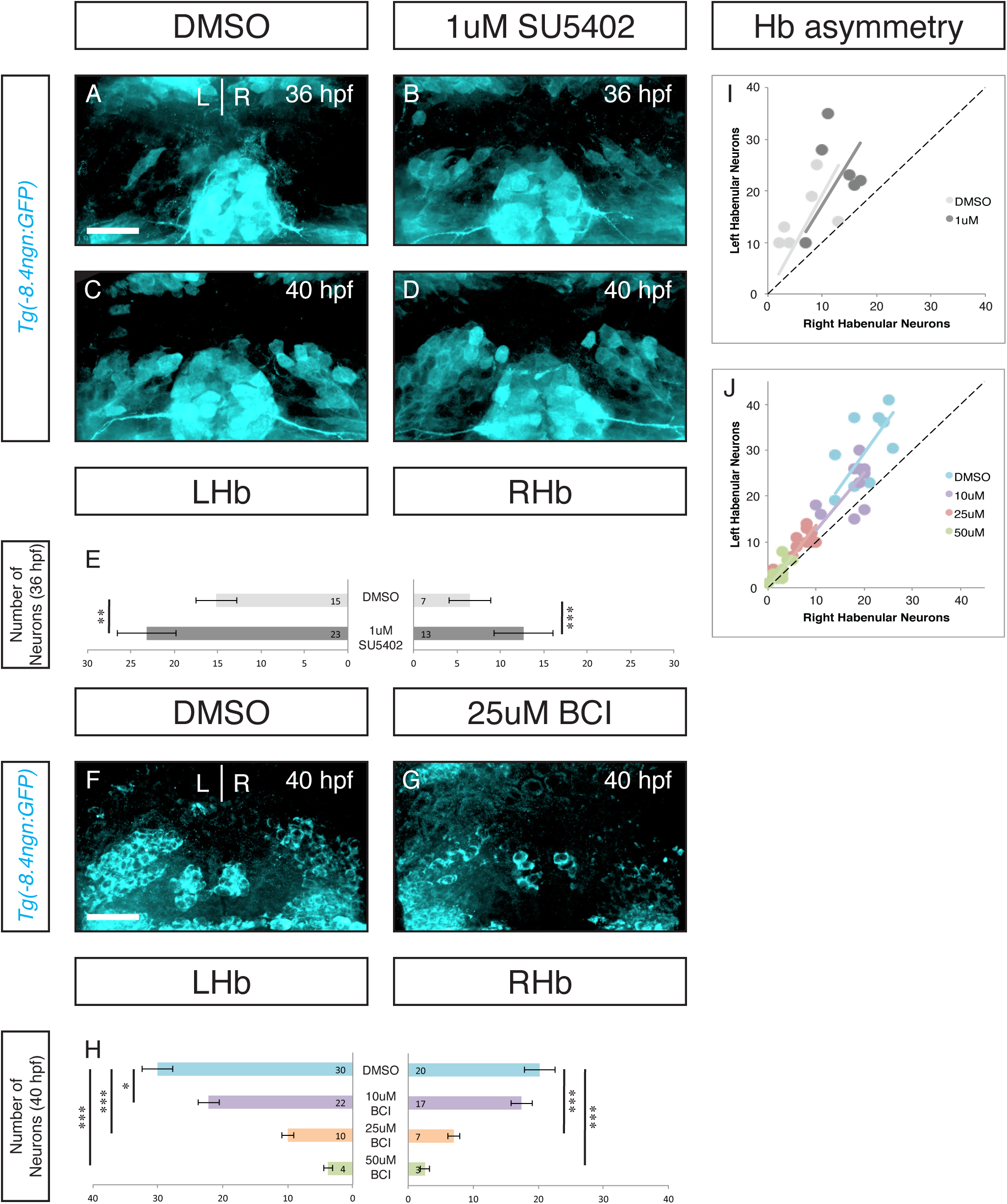
FGF regulates the timing of habenular neurogenesis. Inhibition of FGF signaling by a 1 hour pulse of 1uM SU5402 from 25hpf-26hpf resulted in an increased number of neurons at 36 and 40hpf (A-E). Inhibitor treated embryos retained neurogenic asymmetry (Wilcoxon signed-rank test: for SU5402 p<0.01 for both groups; for BCI p,0.01 for 10uM and p<0.005 for the three remaining groups). *p<0.05, **p<0.01, ***p<0.005. Scale bars are 50uM.

High levels of FGF signaling have previously been associated with progenitor maintenance (Garcia-Maya et al., 2006; Lahti et al., 2010). Given the asymmetric activity of FGF signaling 10 hours before the appearance of neurons, and its sustained activity on the right where neurogenesis begins later, we hypothesized that FGF signaling in habenular progenitors regulates the timing of neurogenesis.

To track neurogenesis we use both transgenic fish with a *neurogenin1* promoter driving GFP (*Tg(-8.4neurog1:GFP)*) and antibody labeling for HuC (a ribosome binding protein that marks post-mitotic neurons). Knowing that *severe* attenuation of FGF inhibits dHb development, we hypothesized that partial reduction of FGF signaling may separate FGFs role in dHb cell survival from a possible role in driving differentiation. To address this we employed a small molecule approach seeking to reduce FGF signaling in a graded fashion. SU5402 is an FGF receptor antagonist. At doses of 12uM, SU5402 has a maximal effect on habenular development phenocopying the *fgf8* null mutants (Data not shown). Using 1uM doses of SU5402 and a short pulse of treatment (1 hour from 25hpf-26hpf), we identified a treatment regimen that had no effect on the number of neurons at 48 hpf (Data not shown). Using this ‘sub-maximal’ dose of FGF antagonist, we measured the appearance of neurons at 36hpf and 40hpf using *Tg(-8.4neurog1:GFP)* zebrafish.

Sub-maximal inhibition of FGF signaling resulted in a premature appearance of neurons at the earliest stages of habenular neurogenesis (Figure 2A-E). Both the left and right habenula show significant increases in GFP+ cells (p<0.01 and p<0.005 respectively, Figure 2E). Interestingly, habenular asymmetry is maintained despite premature neurogenesis (Figure 2I, p<0.01). Thus, sub-maximal inhibition of the FGF signaling pathway led to premature habenular neurogenesis. This is consistent with a model where the level of FGF signaling acts as neurogenic switch, as FGF signaling levels drop below a certain threshold NPs begin to make neurogenic divisions.

A prediction of this model is that if high levels of FGF activity are sustained, there should be a delay in the onset of neurogenesis. BCI is an allosteric inhibitor of Dusp6, an FGF feedback inhibitor. Dusp6 inhibition leads to increased FGF activity, but the increase will not exceed physiologic levels. As Dusp6 is a feedback inhibitor, treating embryos from 25hpf-26hpf in increasing doses of BCI resulted in a dose-defendant delay in habenular neurogenesis (Figure 2F-H). Again, the temporal shift in neurogenesis had no effect on the asymmetry of neurogenesis (Figure 2J, p<0.005). Therefore, tuning FGF signaling up and down is sufficient to regulate the timing of habenular neurogenesis. We conclude that FGF activity acts as a gating mechanism for the timing of habenular neurogenesis.

### FGF represses cyclin-dependent kinase inhibitor, *kip2*, during habenular development

So far we have demonstrated that FGF signaling is asymmetrically deployed to regulate the timing of neurogenesis. But how does FGF regulate the switch from proliferative to neurogenic divisions in NPs? FGF signaling often targets cell cycle regulators and in neural tissues is known to repress Cyclin-dependent kinase inhibitors (CDKIs, Frederick & Wood 2004). This prompted our investigation of the expression of two CDKIs, *kip1/2* in the developing habenula. While *kip1* is not expressed in the habenula (data not shown), *kip2* shows robust habenular expression from 36hpf to 48hpf (Figure 3A, B, C). However, its expression is present at high levels in neighboring domains as well obscuring the details of the its habenular expression. To enhance our analysis we employed the transgenic line, *Tg(dbx1b:GFP)*, which marks habenula and not immediately neighboring tissue. Using this marker, we were able to isolate dHb expression for further analysis. Interestingly, *kip2* expression seems to progress through the left and right habenula in offset waves. Expression in the left habenula reaches a peak between 36hpf and 40hpf before dropping off by 48hpf (Figure 3A’, B’, C’). In the right habenula expression continues to accumulate from 36-48hpf. At 36hpf and 40hpf there is significantly more *kip2* expression in the left habenula (Figure3 A”&B”). These data show that *kip2* is asymmetrically expressed during the onset of habenular neurogenesis and therefore might be target of habenular FGF signaling. Indeed, *kip2* is significantly upregulated in *fgf8^x15/x15^* mutants (Figure 3F&G). Thus we propose that early high levels of FGF inhibit *kip2* expression allowing NPs to retain their progenitor status, but as FGF levels decline, *kip2* is upregulated and drives progenitor daughters to exit the cell cycle and begin differentiation. This processes happens earlier in the left habenula.

**Figure 3:**
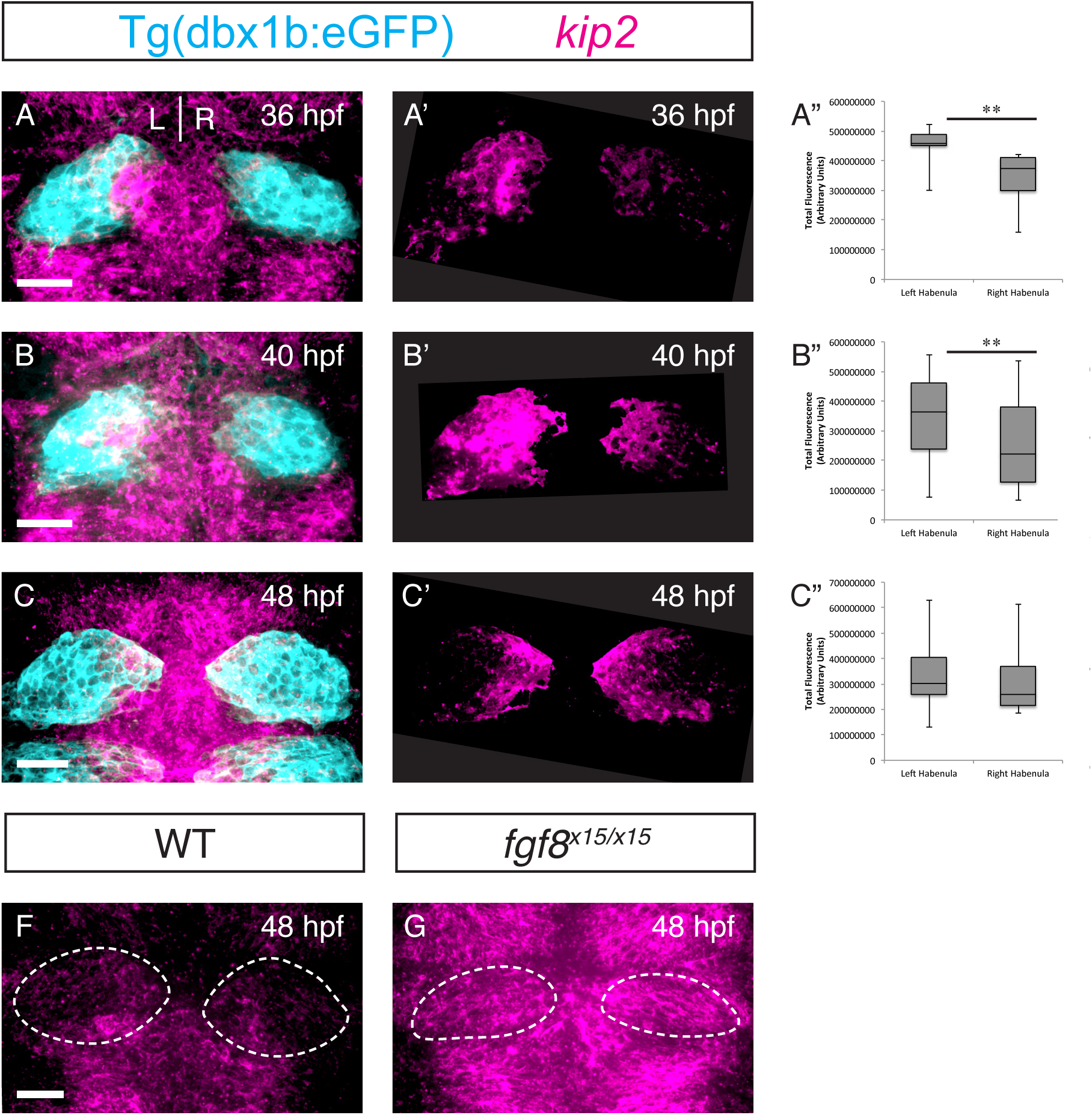
*kip2* appears in asymmetric pulses and is regulated by FGF. *kip2* and a transgene marking the habenula are coexpressed (A, B, C). A pulse of *kip2* expression appears between 36hpf and 48hpf. The pulse peaks on the left at 40hpf and is still increasing on the right by 48hpf (A’, B’, C’). *kip2* expression is leftward biased at 36hpf and 40hpf (A”, B”; Wilcoxon signed-rank test: 36hpf and 40hpf p<0.025). By 48hpf the levels of *kip2* are no longer significantly different (C”). **p<0.01. Scale bars are 50uM.

In addition to inhibiting cell cycle exit, FGF signaling is also known to directly promote NP maintenance (Lahti et al., 2010). *Her6* is a member of *hes/her* family of pro-progenitor transcription factors. *Her6* is a negative regulator of *ngn1* has been shown to regulate the timing of neurogenesis and neuron cell fate in the adjacent thalamus (Scholpp et al., 2009; Yoshizawa et al., 2011). As well, *her6* expression in the habenula has been reported to have a right-sided bias (Aizawa et al., 2007). Using the same methods as our *kip2* analysis, we observed a complementary expression pattern. After early, bilaterally high, levels of expression, *her6* expression declines on the left before the right (Supplemental Figure 2 A-A’, B-B’). Expression is significantly lower in the left habenula just before neurogenesis begins (Supplemental Figure 2A”). While *her*/*hes* transcription factors are canonically downstream of the Notch signaling pathway, there are conflicting reports concerning weather *her6* is or is not regulated in a Notch-independent manner (Aizawa et al., 2007; Hans et al., 2004; Scholpp et al., 2009). This confusion and the correlation of FGF and *her6* asymmetry raise the possibility that FGF may regulate progenitor maintenance via *her6*. Together the *kip2* and *her6* data strongly suggest that FGF gates habenular neurogenesis by inhibiting CDKIs and possibly by maintaining expression of pro-progenitor factors.

### Nodal signaling leads to early down regulation of FGF signaling in the left habenula

FGF signaling appears to be the center of neurogenic cassette that is asymmetrically regulated in the developing habenula. What is source of the asymmetric regulation of FGF activity? Nodal signaling, specifically its effector in the zebrafish CNS, *cyclops*, is know to establish the asymmetric timing of habenular neurogenesis (Roussigné et al., 2009). We wanted to determine if left-sided Nodal activity drove the asymmetry in FGF signaling we observed. In optical sections, the Nodal target *lefty1* is expressed in the same plane as the asymmetric FGF signal at 26hpf (Figure 4A&B). As well, *lefty1* expression colocalizes with the habenular marker *dbx1b* (Figure 4C&D). Thus nodal signaling components are active in the habenula during asymmetric FGF activity.

**Figure 4:**
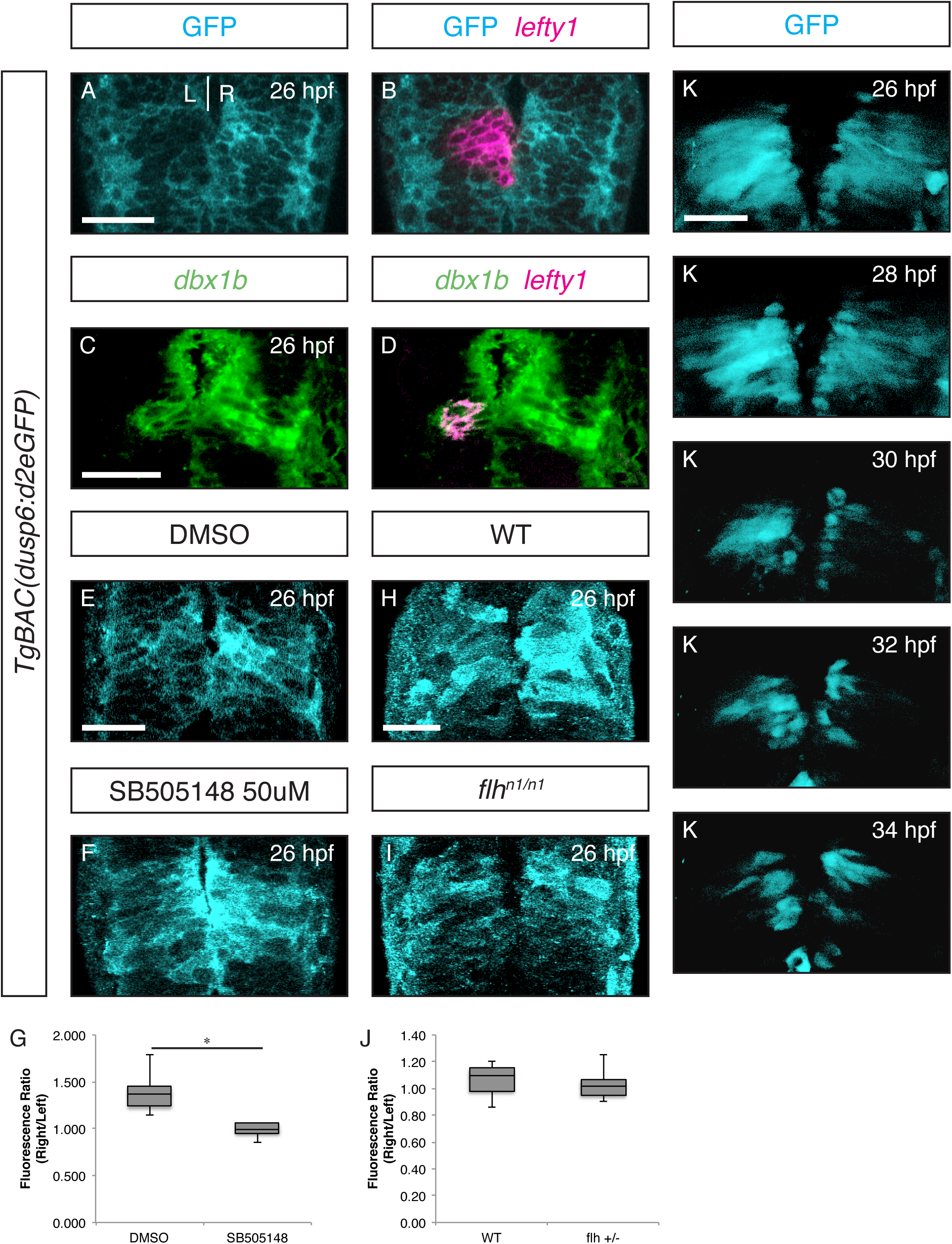
Nodal regulates FGF signaling in the developing habenulae. The Nodal signaling component *lefty1* is expressed in the plane of asymmetric FGF activity (A-B). As well, *lefty1* is co-expressed with the habenular marker *dbx1b* on the left (C-D). Inhibition of Nodal by the small molecule or instantiation of bilateral Nodal in the *flh* mutant led to a symmetrization of FGF signaling activity (E-J). This was significant in the case of SB505124 treatment (G-J, Wilcoxon signed-rank test: for SB505124 p<0.025). As well, in vivo time-lapse imaging show that symmetrization of FGF activity persisted through 34hpf after drug treatment (K-O). *p<0.05, **p<0.01, ***p<0.005. Scale bars are 50uM.

**Figure 5:**
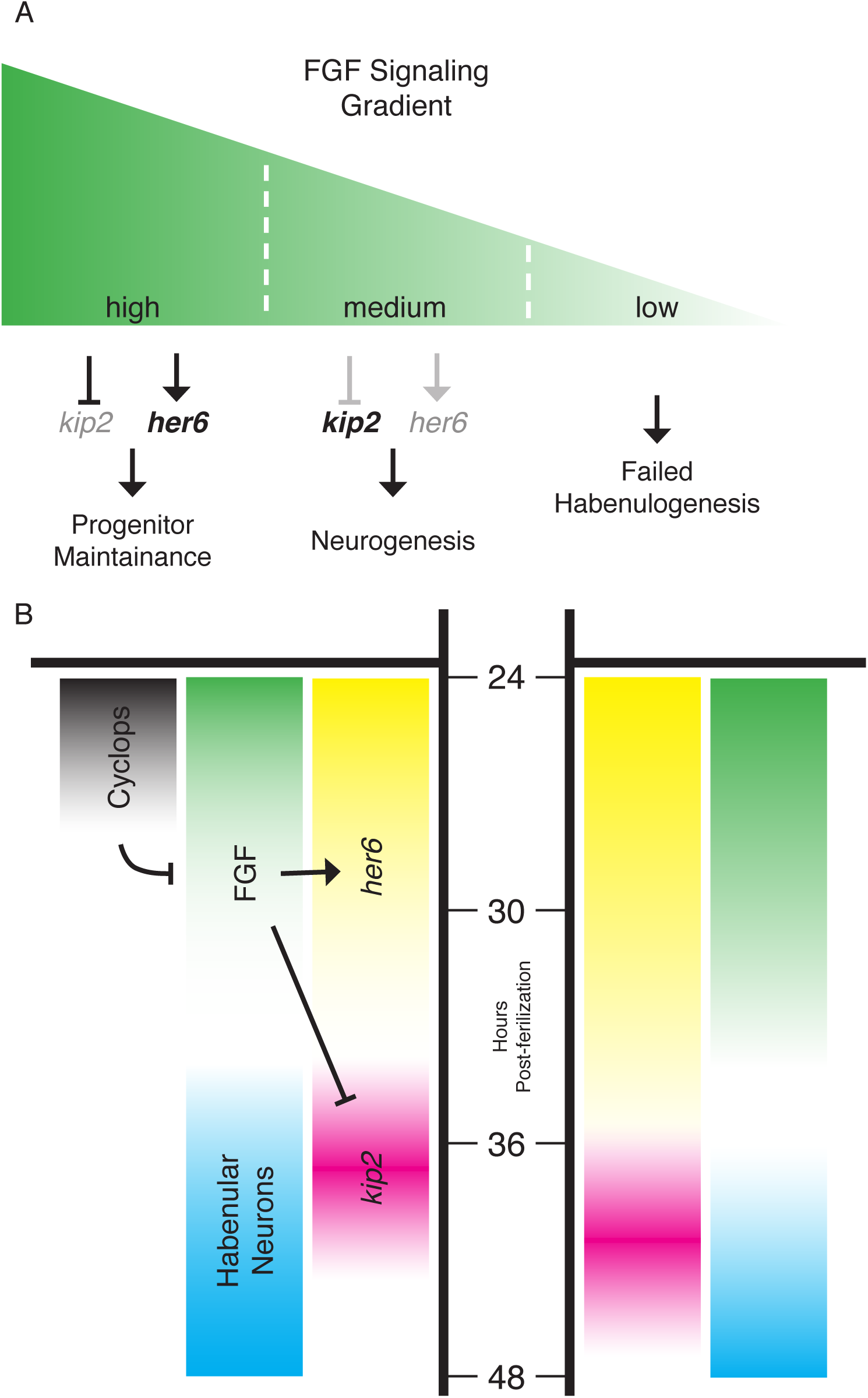
Levels of FGF signaling regulate the timing of neurogenesis in the habenula. Nodal inhibits FGF signaling on the left resulting in earlier neurogenesis. We propose a model where early high levels of FGF inhibit the CDKI *kip2* and promote pro-progenitor factors such as *her6*. This maintains habenula NPs in a progenitor state.As levels of FGF signaling drop, kip2 is derepressed and her6 expression decreases. This allows NPs to begin neurogenic divisions where neuronal daughters exit the cell cycle and turn on pro-neural genes (HuC, ngn1). Drastic reduction of FGF signaling (fgf8x15/x15 null mutant) leads to failed habenulogenesis sue to decreased proliferation, increased apoptosis and failed differentiation (A). FGF is a key regulator of a neurogenic cassette (along with Notch). Asymmetric habenular neurogenesis is achieved by left-sided Nodal inhibition of FGF signaling. This initiates the decline of FGF signaling earlier on the left (B).

To directly test if Nodal downregulates FGF signaling in the left habenula, we treated embryos with a Nodal receptor antagonist SB505124 that targets the ALK3/4 receptors selectively. We treated embryos with 50uM SB505124 from 16-24hpf, a time range chosen to avoid early requirements for Nodal in lateral plate mesoderm patterning and neural plate induction. SB505124 treatment led to a symmetrization of FGF signaling activity at 26hpf (Figure 4E-G). Indeed there is an increase in FGF activity on the left in the absence of Nodal signaling (Figure 4E&F). Due to midline defects, Nodal signaling is activated bilaterally in *floating head (flh)* mutants. So we tested what effect there might be on FGF activity with bilateral Nodal signaling. In this condition, FGF signaling also becomes symmetric, but with a decrease in FGF activity on the right (Figure 4H-J). Together, these experiments show that asymmetric FGF signaling is established by left-sided Nodal inhibition of FGF activity. To our knowledge this is the first reported regulation of FGF by the Nodal signaling pathway.

Several habenular asymmetries in the zebrafish are established by the leftward migration of the small accessory organ, the parapineal. Asymmetric habenular neurogenesis is known to be parapineal-independent (Roussigné et al., 2009). In *tbx2b^c144/c144^* mutants, there is failure to form a left-sided parapineal. To determine if asymmetric FGF signaling was parapineal-dependent, we analyzed FGF signaling activity in *tbx2b ^c144/c144^* mutants and found no change in asymmetry of FGF signaling (Supplemental Figure 3). Thus, asymmetric FGF activity regulates the asymmetric onset of habenular neurogenesis. This neurogenic gate is rendered asymmetric by left-sided Nodal signaling in a parapineal-independent manner.

## DISCUSSION

Here we report an FGF regulatory cassette that determines the timing of neurogenesis in the habenular nuclei. Early in habenular development previously established FGF signaling begins to diminish. This derepresses the cell-cycle dependent kinase inhibitor, *kip2* (and may be the cause of the downregulation of the pro-progenitor transcription factor *her6*). The pulsatile expression of *kip2* helps drive habenular neural progenitors to begin neurogenic divisions leading to the appearance of the first habenular neurons. This neurogenic program proceeds asymmetrically with neurons maturing in the left habenula several hours before they appear in the right. This syncopation of neurogenesis is a result of Nodal signaling acting in the left habenula to attenuate FGF signaling earlier on that side. Thus, FGF serves as a crucial point of signal integration between L/R patterning driven by Nodal and spatiotemporal patterning to define habenular cell fate.

Several key questions follow from this exciting discovery. The Notch signaling pathway is crucial to habenular neurogenesis and its manipulation can also alter the timing of neurogenesis (Aizawa et al., 2005). However, no clear asymmetries in Notch signaling have been reported. Thus it is unknown how Notch-mediated habenular neurogenesis is rendered asymmetric. How are asymmetric FGF signaling and Notch signaling integrated? Given the contradictory literature on how *her6* is regulated, it may be the case that Notch and FGF signaling converge at *her6*. Aizawa et al. (2005) also demonstrated a correlation between the timing (early vs late) of habenular neurogenesis and the fate of a neuron. They observe that early-born neurons take on a lateral subnucleus fate, while late-born-neurons take on a medial subnucleus fate. If FGF regulates the timing of neurogenesis, do perturbations of FGF-mediated neurogenesis have an effect on cell fate? Indeed, preliminary analysis shows that precocious neurogenesis (from FGF attenuation with SU5402 treatment) results in an increase in lateral subnucleus (Kctd12.1+) neurons (Supplemental Figure 4 A-D). It will be exciting to see if BCI-treated embryos show a decrease in this same population and an increase in medial subnucleus neurons.

By studying neurogenesis in simple model systems we have uncovered exciting new targets and pathways regulating the NP switch from proliferative to neurogenic cell fate decisions as well as validating a small molecule approach to manipulating those targets and pathways. To our knowledge this is the first report of Nodal regulation of FGF signaling – two signaling pathways central to stem cell maintenance and differentiation (Sui et al., 2013). Still, the molecular mechanisms connecting the two pathways are unclear. Transcriptome analysis has shown Nodal to be upstream of *dusp4* in the developing lateral plate mesoderm, a member of dual-specificity phosphatase family capable of down regulating FGF signaling (Brown et al., 2008). However, currently we found no evidence for asymmetric activation of *dusp4* during habenulogenesis (data not shown). In addition to the Nodal regulation or FGF activity, our use of small molecules *in vivo* to tune endogenous signaling pathways and regulate neurogenesis contributes to a growing body of work defining how small molecules can be used to determine cell fate (Lu and Atala, 2014).

This is a very exciting milieu of tools and targets to apply to questions of basic neurodevelopment as well as regenerative medicine. Therapeutic application of neurons is coming closer to a clinical reality. There is a large amount of work to be done to develop stem cell protocols that deliver high yields of specific cell types, and treatments to ensure their survival and successful integration into the nervous system (Anderson and Vanderhaeghen, 2014; Deidda et al., 2014; Southwell et al., 2014). The dHb can provide a wonderful model to explore for very real contributions to the next generation of clinical stem cell interventions.

## Figure Legends

**Supplementary Figure 1:**
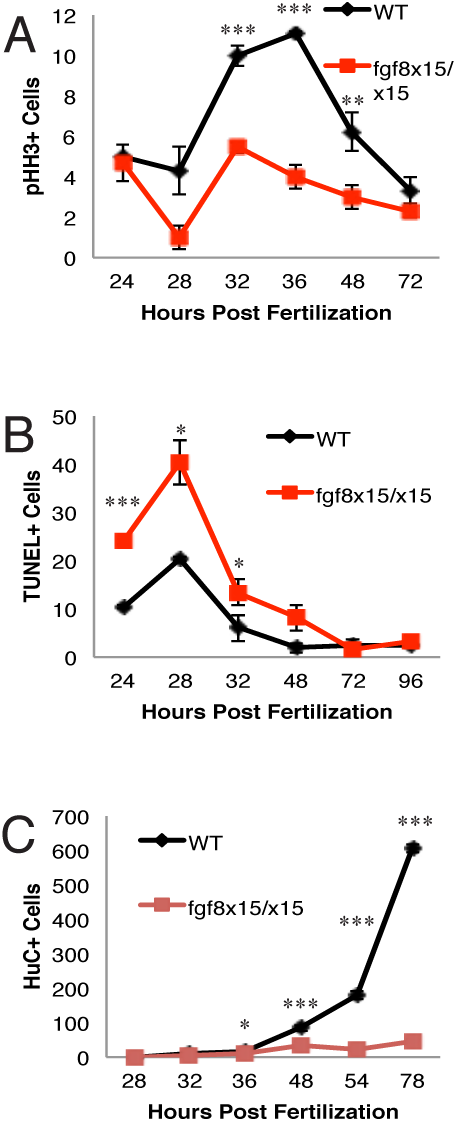
*fgf8* mutants show reduced proliferation, increased cell death and failed differentiation. *fgf8* mutants show signif-cantly fewer pHH3+ cells between 32 and 48hpf (A). They also show more TUNEL+ cells between 24 and 32hpf (B). Finally, the remaining cell fail to differentiate into neurons (C). *p<0.05, **p<0.01, ***p<0.005.

**Supplementary Figure 2:**
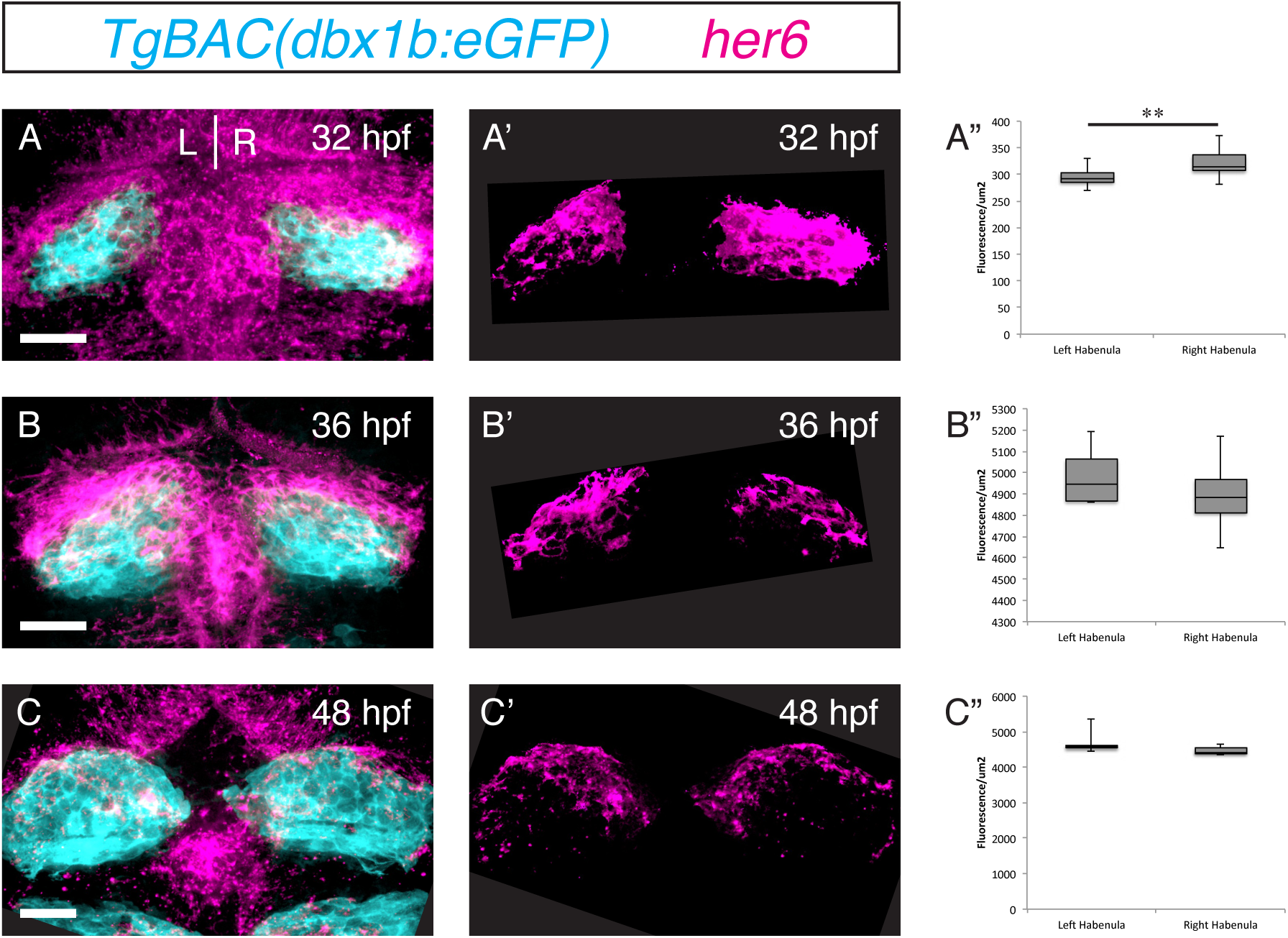
*her6* expression is present at high levels on the right early in habenular development. *her6* is expressed at higher levels in the right ha-benula at 32hpf (A-A”). By 40 and 48hpf expression levels have reduced and become symmetric (B-C”). *p<0.01. Scale bars are 50uM.

**Supplementary Figure 3:**
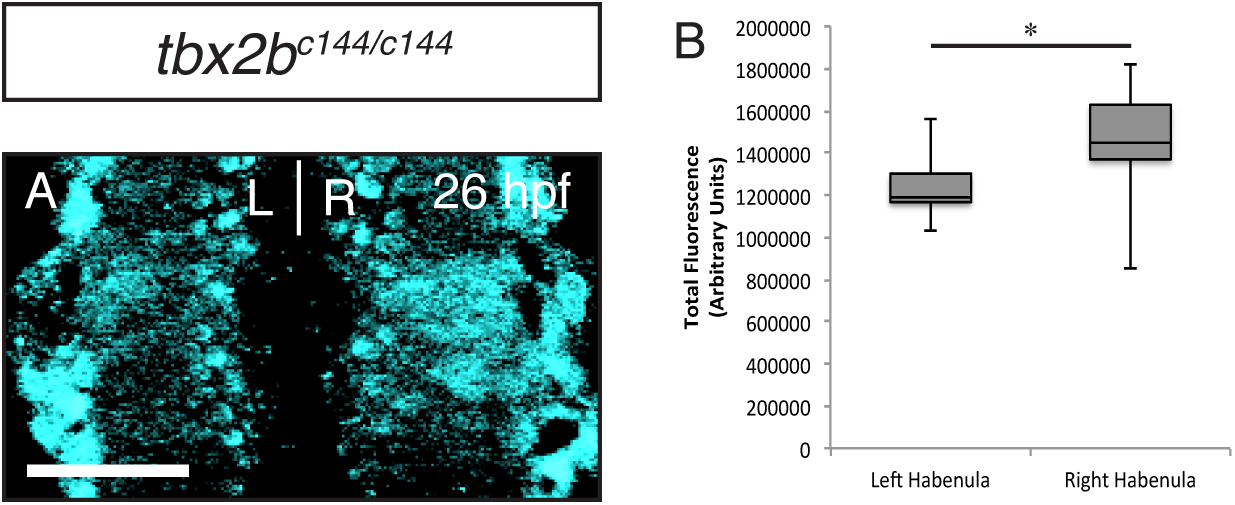
Right-biased FGF activity is independant of parapineal development. *tbx2b* mutants retain right-biased FGF activity at 26hpf (A-B). *p<0.05. Scale bar is 50uM.

**Supplementary Figure 4:**
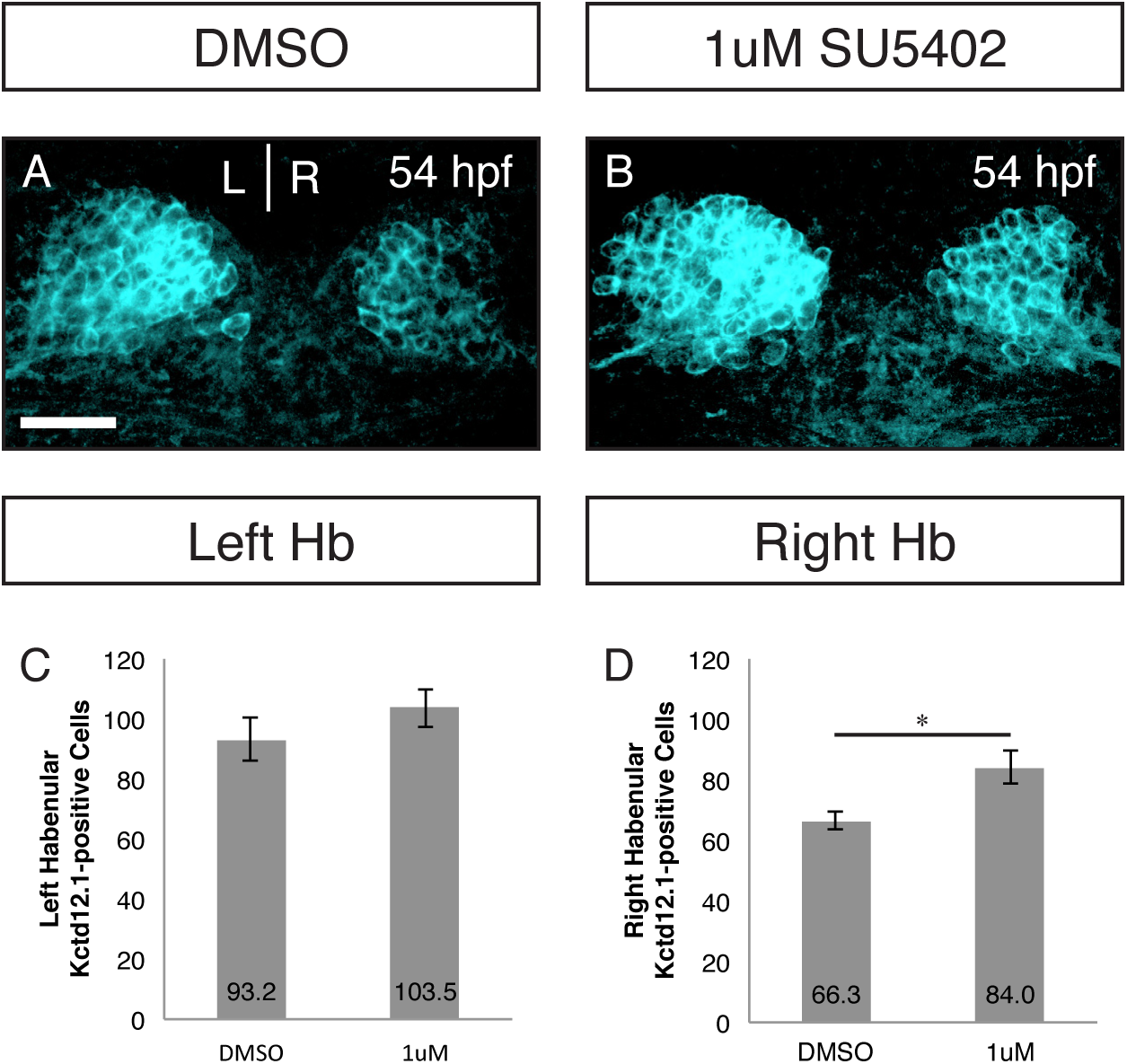
Timing of neurogenesis impacts habenular cell fate. SU5402 treatment reults in an increase in Kctd12.1 neruons by 54hpf in the left habenulae and to a signifcant degree in the right habenula (A-D). *p<0.01. Scale bar is 50uM.

